# DJ-1/PARK7 Determines miRNA Network Plasticity During Genotoxic Stress

**DOI:** 10.64898/2026.08.10.744036

**Authors:** Keren Zohar, Michal Linial

**Affiliations:** Department of Biological Chemistry, Institute of Life Sciences, The Hebrew University of Jerusalem, Jerusalem, Israel

**Keywords:** RNA-seq, regulated cell death, lncRNA, oxidation stress, X-ray, ribosome stability, miRNAs, siRNA, Parkinson disease, Functional Genomics

## Abstract

PARK7 (DJ-1) is a redox-sensitive stress-response protein that supports cellular adaptation, but its role in post-transcriptional responses to genotoxic stress remains unclear. We investigated whether DJ-1 abundance determines the miRNA response to X-ray–induced DNA damage. Integrated mRNA-seq and small RNA-seq across DJ-1 states in HEK293 cells revealed a striking divergence in miRNA plasticity. DJ-1 depletion by siRNA produced minimal miRNA remodeling, with only 34 (6.3%) miRNAs differentially expressed after irradiation. In contrast, elevated DJ-1 markedly increased miRNA plasticity: irradiation altered ∼37% of detectable miRNAs, accounting for ∼90% of miRNA reads, and extensively redistributed the miRNA pool. DJ-1 overexpression was also associated with remodeling of the miRNA regulatory machinery, particularly components involved in miRNA sorting and stability, suggesting feedback regulation of the miRNA pool. Comparison of precursor and mature species revealed substantial uncoupling between transcription and mature miRNA abundance, implicating regulation at the levels of processing, maturation, or stability. Radiation-responsive coding genes in DJ-1-overexpressing cells were relatively depleted of miRNA binding sites, supporting preferential regulation of upstream regulatory nodes rather than the bulk transcriptome. Together, these findings identify DJ-1 as a determinant of post-transcriptional signaling plasticity, enabling dynamic remodeling of the miRNA regulatory state in response to genotoxic stress.

**Graphical Abstract:** 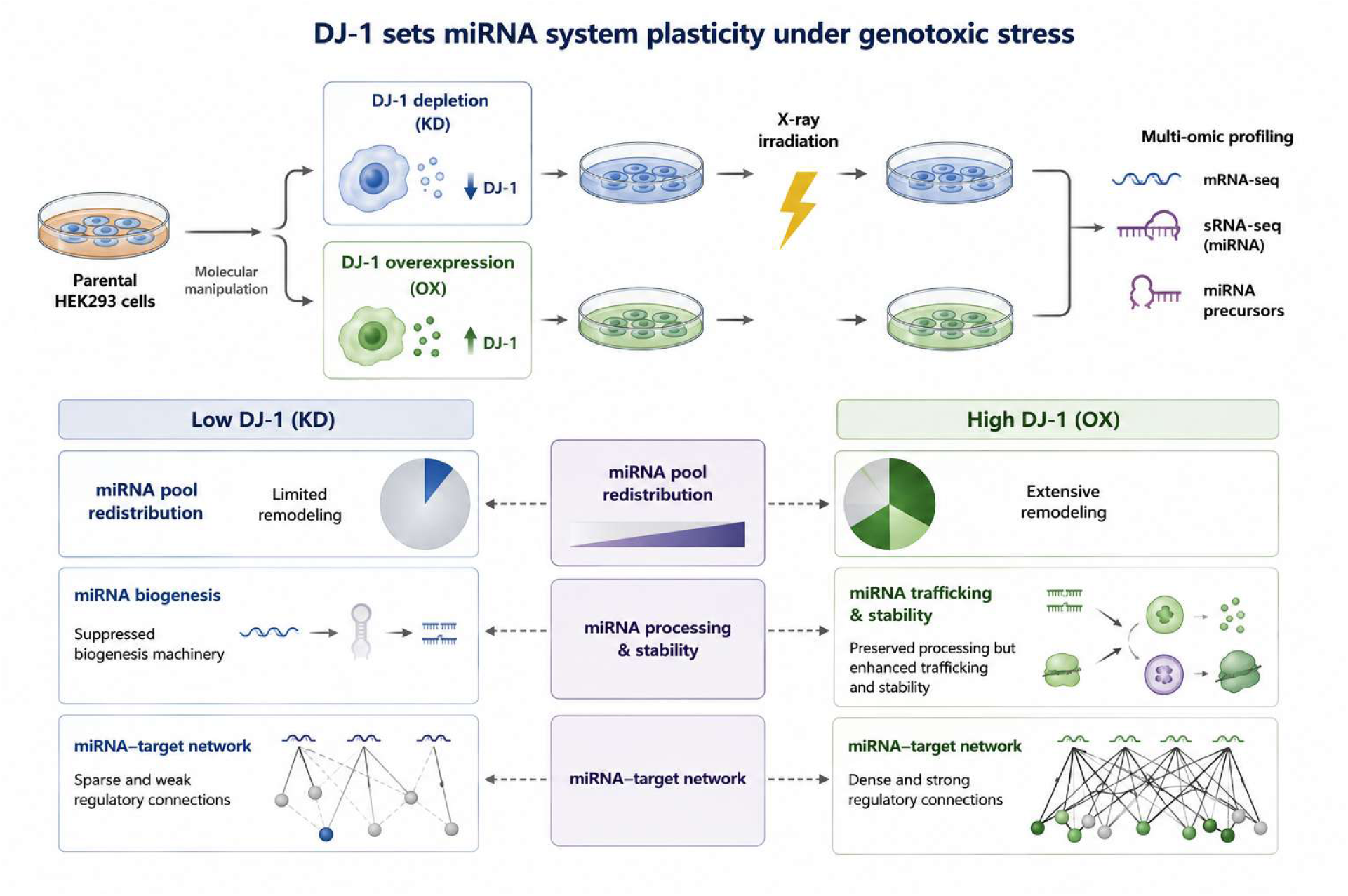

## Introduction

PARK7 (DJ-1) is a redox-sensitive stress response protein essential for oxidative defense and mitochondrial homeostasis, yet its role in post-transcriptional regulation during genotoxic stress remains unclear (Baulac et al, 2009; Kahle et al, 2009). PARK7, also known as DJ-1, is a highly conserved multifunctional protein that promotes cellular resilience to diverse stressors, including oxidative stress and genotoxic insult (Higgins et al, 2010; Lev et al, 2008). DJ-1 functions as an antioxidant and redox sensor, modulates transcriptional networks, contributes to mitochondrial quality control, and regulates signaling pathways that govern cell survival and metabolism (Ariga et al, 2013; Dolgacheva et al, 2019; Wilson, 2011). A critical determinant of DJ-1 activity is its highly reactive cysteine residue at position 106 (Cys106), which serves as a redox sensor essential for stress-responsive functions. Mutation of Cys106 (C106S) disrupts DJ-1 dimerization (Ariga et al, 2013) and impairs its nuclear functions (Bhattacharyya et al, 2022). Pathogenic mutations in PARK7 are linked to familial Parkinson’s disease, underscoring the importance of DJ-1 in protecting neurons from reactive oxygen species (ROS) and maintaining genome stability (Oh et al, 2018; Pfeifer, 2024; Xiao et al, 2022). Consistently, in vivo studies show that DJ-1 deficiency exacerbates tissue damage under oxidative stress, including cardiac injury, highlighting its broader role in stress adaptation beyond neurodegeneration (Kahle et al, 2009).

Although DJ-1’s roles in oxidative stress responses and mitochondrial regulation are well established, its involvement in post-transcriptional gene control remains poorly defined. In parallel, DJ-1 was shown to link to noncoding RNA (ncRNA) regulation layers (Zohar et al, 2022). Specifically, DJ-1 (PARK7) likely regulates genotoxic stress responses and influence the expression of stress-induced lncRNAs that control major cell decisions (e.g., cell-cycle progression, ribosome biogenesis, transcriptional attenuation). Such lncRNAs are established regulators of chromatin architecture and RNA polymerase II activity (Gámez-Valero et al, 2020). In parallel, by preserving proteostasis under oxidative stress, DJ-1 may stabilize RNA-binding proteins and nucleolar components, thereby indirectly affecting snoRNA-guided rRNA modification and ribosome remodeling, processes central to translational control during DNA damage (Chen et al, 2024; Ni et al, 2022). Together, these mechanisms position DJ-1 as an upstream regulator of ncRNA-dependent translational buffering.

Across the animal kingdom, particularly in mammals, the layer of microRNAs (miRNAs) has been extensively studied in both health and disease (Gebert & MacRae, 2019). In the context of cancer, differentially expressed miRNAs are considered sensitive indicators of cellular state changes and disease progression (Pardini et al, 2018). In humans, more than 2,500 mature miRNAs, encoded by approximately 1,900 genes, have been annotated, and they predominantly function to repress gene expression at the post-transcriptional level (Kozomara et al, 2019). miRNAs are increasingly recognized as key regulators of cellular stress responses (Babar et al, 2008; Emde & Hornstein, 2014). Studies of UV-induced DNA damage have demonstrated that cell-cycle regulation and apoptosis are controlled not only transcriptionally but also by miRNAs (Bueno et al, 2008; Wang & Lee, 2009), accompanied by rapid miRNA expression changes, and relocalization of AGO2 to stress granules (Olejniczak et al, 2018; Pothof et al, 2009). In cancer and other stress contexts, miRNAs dynamically reshape protein output by altering biogenesis, processing, and target selection in response to DNA damage and environmental cues (Olejniczak et al, 2018; van Jaarsveld et al, 2014). Notably, specific DNA damage-responsive miRNAs regulate cell-cycle checkpoints (Hu & Gatti, 2011), DNA repair pathways, and survival decisions following genotoxic treatments, including ionizing radiation (Hattori et al, 2014; Mao et al, 2014).

Emerging evidence suggests direct links between DJ-1 and miRNA networks. For example, DJ-1 regulates miR-221 in neuronal models, promoting cell survival through repression of apoptotic factors, an effect lost in pathogenic DJ-1 mutants (Oh et al, 2018). In addition, DJ-1 has been implicated in DNA damage repair through interactions with core repair enzymes, linking it to the DNA damage response (DDR) at the level of genome maintenance (Wang et al, 2023). While X-ray irradiation robustly activates the DDR to halt replication and facilitate repair, it remains unclear whether DJ-1 directly contributes to this response. Persistent DNA damage, particularly in neurons with compromised repair capacity, is thought to drive genome instability, mitochondrial dysfunction, and neurodegeneration in Parkinson’s disease (Gonzalez-Hunt & Sanders, 2021; Vazquez-Villasenor et al, 2021). Moreover, whether DJ-1 coordinates global miRNA reprogramming in response to genotoxic stress, and how this impacts translational control and stress adaptation, remains unresolved.

To address this gap, we previously used a controlled experimental system in which DJ-1 expression was manipulated to examine the effects of X-ray–induced DNA damage on mRNA transcriptome dynamics (Zohar et al, 2025). We found that DJ-1 overexpression in following X-ray licenses a stress-dependent reprogramming that restrains protein synthesis at cytoplasmic and mitochondrial ribosomes, while preserving adaptive stress programs and reshaping lncRNA expression (Zohar et al, 2025). In this study we tested the contribution of miRNAs in cellular homeostasis and stress response under genotoxic stress. Using high-throughput small noncoding RNA-seq (sncRNA-seq), we focused on hundreds of miRNAs and inferred their impact of pathway analyses. Together, the findings from whole transcriptome levels (mRNAs, lncRNAs and miRNAs) position DJ-1 as a sensor and integrator of DNA damage signaling. We found that excess of DJ-1 permits miRNA remodeling under genotoxic stress. Thus, we extending the canonical role of DJ-1 role from oxidative stress defense to post-transcriptional control under genotoxic stress.

## Materials and Methods

### Manipulating DJ-1 levels in cells

Human embryonic kidney 293 cells (HEK293, ATCC) were cultured in 6-well plates with 70–80% confluence, at 37°C and 5% CO₂ in Dulbecco’s modified Eagle’s medium (DMEM) high-glucose supplemented with 10% fetal calf serum (FCS) as described before (Zohar et al, 2022). HEK293 cells were plated in a 10-cm plate at an initial density of 5×10⁶ cells per plate one day prior to their transfection. The pCMV3 vector encoding canonical DJ-1, fused to tandem Strep-tags at its C-terminus, was obtained from Sino Biological. The Strep-tag peptide exhibits a high intrinsic affinity for Strep-Tactin, an engineered streptavidin. The expression plasmid (DJ-1 or empty pCMV3 vector) and PEI (260008-5; Polysciences) were separately diluted in Opti-MEM I (31985-047; Gibco), then mixed and incubated at room temperature for 25 min prior to addition to the cells. The mixture was added dropwise on the cell culture. After 18 h, cells were exposed to X-ray radiation (parameters: 12.5 mA, 320 kV, 10 Gy) and total and small RNAs were extracted six hours later. Notably, the endogenous DJ-1 levels remained unchanged, demonstrating that overexpression specifically increased exogenous DJ-1 without altering the native DJ-1 protein (Zohar et al, 2025).

For DJ-1 knockout (KD) by siRNA protocol, cells were cultured in a 6-well plate at 20–30% confluence (3 × 10^4^ per cm^2^). For transfection, we used Lipofectamine 2000 (Lipo2000; Invitrogen, Cat #11668019, Carlsbad, CA, USA) at 40–60% confluence, following the manufacturer’s instructions. The esiRNA MISSION system, (Sigma-Aldrich, Burlington, MA, USA) was used with a control of esiRNA-RULC, as a negative control (marked as RLUC), alongside the specific PARK7 esiRNA (# EHU113961). After 18 h, cells were exposed to X-ray radiation (parameters: 12.5 mA, 320 kV, 10 Gy), and 6 h later, total RNA was extracted and was prepared for sncRNA-seq and RNA-seq libraries (for details see (Zohar et al, 2025)).

### RNA extraction and library preparation

Total RNA was extracted using the RNeasy Plus Universal Mini Kit (QIAGEN, Cat #73404, Redwood City, CA, USA) according to the manufacturer’s protocol. One microgram of total RNA was used for poly(A) selection to enrich for mRNAs. Libraries were prepared with the KAPA Stranded mRNA-Seq Kit following the manufacturer’s instructions and sequenced on an Illumina NextSeq 500 platform to generate 85 bp single-end reads, yielding 25–30 million mapped reads per sample.

Small noncoding RNA-seq (sncRNA-seq) was prepared from total RNA isolation using standard column-based methods and assessed for integrity and purity. For each sample, 100 ng of RNA (<200 bp) was used as input for miRNA library preparation. Small RNAs were selectively ligated to adapters using a sequential ligation strategy. Briefly, a 3′ RNA adapter was first ligated to the hydroxyl group at the 3′ end of small RNAs using T4 RNA ligase, followed by ligation of a 5′ RNA adapter to the 5′ phosphate. Adapter-ligated small RNAs were reverse-transcribed using adapter-specific primers to generate cDNA. The resulting cDNA was amplified by PCR using Illumina-compatible indexed primers to enable sample multiplexing. PCR cycle numbers were minimized to reduce amplification bias. Size selection was performed to enrich for fragments corresponding to mature miRNAs (75 bp) using either gel-based purification. Purified libraries were assessed for size distribution using a Bioanalyzer system and quantified. Equimolar amounts of indexed libraries were pooled and sequenced on an Illumina platform using single-end reads (75 bp). Base calling and demultiplexing were performed using standard Illumina software.

### Differential expression of miRNAs (DEMs)

Next-generation sequencing (NGS) data underwent quality control using FastQC (v0.11.9), followed by preprocessing with Trimmomatic (v0.32). Adapter trimming was performed using cutadapt, removing the 3′ adapter sequences (AGATCGGAAGAGCACACGTCTGAACTCCAGTCAC and CGATC) and retaining reads with a minimum length threshold of 18 nucleotides. Processed reads were aligned to the reference genome (GRCh38) using miRDeep2. The quantification of miRNAs was performed using miRDeep2 with miRBase v22 annotations. Lowly expressed genes were filtered out using a threshold of at least two counts per million in three samples. Raw miRNA read counts were normalized using the trimmed mean of M-values (TMM) method implemented in edgeR (version 3.36.0), which corrects for compositional differences between libraries.

Differential expression (DE) analysis was performed for all experimental groups. Genes with FDR <0.05 and an absolute log fold change (FC) ≥|0.5| were considered up- or downregulated, respectively, all others genes were considered unchanged. Principal component analysis (PCA) was performed using the R base function prcomp (RStudio v4.1.0). Differential expression analysis was conducted with edgeR (v3.36.0), using trimmed mean of M-values (TMM) normalization. Multiple comparisons were corrected using the false discovery rate (FDR) threshold of 0.05. edgeR was selected for its robustness to deviations from normality. Figures were generated using the ggplot2 R package (v3.3.5).

We quantified miRNAs by read counts (per million reads). We refer to the percentage of the ncRNAs from two normalize libraries (sncRNAs and RNA-seq). These proportions refer to the composition of reads within each respective sequencing library and should not be interpreted as absolute cellular RNA abundance.

### Bioinformatic analysis

We are using the nomenclature of miRNAs to cover different aspects of analysis. Specifically, miRNA gene and precursor transcript refer to the biotype labelled miRNA transcript that was detected in the >200 nt total RNA library. The mature miRNA is the processed ∼22–24 nt species detected in sncRNA-seq library. For the analysis of mature miRNA with respect to the precursor identity, we unify 5p and 3p arms to a single entity.

RNA and protein expression data for PARK7 (DJ-1) were retrieved from the Human Protein Atlas (HPA) database (Digre & Lindskog, 2021), including GTEx tissue profiles and cell line datasets (Consortium, 2020). Experimentally validated miRNA–target interactions were obtained from miRTarBase v9.0 and integrated using miRNet 2.0 for network-based visual analytics of differentially expressed miRNAs (DEMs) (Chang & Xia, 2022). Protein–protein interaction (PPI) networks were constructed using STRING with a high-confidence interaction score (>0.7). Network connectivity excluded gene neighborhood, gene fusion, and co-occurrence evidence (Szklarczyk et al, 2025). Statistical significance of PPI enrichment was assessed using STRING’s built-in enrichment test. Functional enrichment analyses were performed using KEGG, WikiPathways, and over-representation analysis implemented in (Szklarczyk et al, 2025), with false discovery rate (FDR) correction applied for multiple testing. AGO-CLIP-supported miRNA-target interactions were retrieved from ENCORI to quantify independent experimental validation events. Total miRNA binding sites were extracted from TargetScan 8.0 (Agarwal et al, 2015) using broadly conserved binding sites. The number of total and unique miRNA binding sites was calculated per the canonical transcripts (according to Refseq).

## Results

### DJ-1 is stably expressed across cell lines and tissues

To assess whether cellular DJ-1 abundance influences the response to X-ray irradiation, we first examined endogenous DJ-1 expression under basal conditions. Establishing the baseline level of DJ-1 was important for evaluating the DNA damage response (DDR) while minimizing excessive cellular stress and cell death. X-ray irradiation induces DNA double-strand breaks (DSBs) and oxidative stress, including increased reactive oxygen species (ROS), to which DJ-1 contributes through its redox-regulatory functions. Phosphorylation of histone H2AX at Ser139 (γ-H2AX) was used as a transient marker of the local DDR at sites of DSBs (Zohar et al, 2025). Supplementary **Fig. S1** summarizes PARK7 expression across major human tissues and commonly used human cell lines, based on normalized transcript per million (nTPM). Among tissues, skeletal and cardiac muscle, several brain regions, and endocrine glands showed somewhat higher PARK7 expression, consistent with their high metabolic and oxidative demands. Most commonly used human cell lines also expressed substantial levels of PARK7, with an average of approximately 500 nTPM. Consistent with the transcript-level data, DJ-1 protein was abundant across the cancer-related cellular models examined, with an estimated abundance of ∼6.5 mM (Supplementary **Fig. S1**). Together, these observations indicate that DJ-1 is broadly and robustly expressed across tissues and cell types. All subsequent experiments were therefore performed in HEK293 cells, which exhibit typical basal levels of endogenous DJ-1.

### Naïve cells show a minimal miRNA response to X-ray irradiation

We next investigated whether miRNAs contribute to the adaptive response of HEK293 cells to X-ray irradiation. Small non-coding RNA sequencing (sncRNA-seq) was performed across five experimental conditions, including untreated cells (N.T.), cells subjected to DJ-1 knockdown (KD) with the corresponding non-specific siRNA control (RLUC), and the effects of 10 Gy X-ray irradiation under these conditions. miRNA profiles were assessed 6 h after irradiation (Supplementary **Tables S1**). Calibration of the impact on miRNA machinery and targets and following X-ray irradiation across the first 12 h was used to select the 6 h time interval (Rzeszowska-Wolny et al, 2022).

Because X-ray-induced DSBs are accompanied by oxidative stress, we asked whether the strikingly robust cellular response to irradiation was associated with changes in miRNA expression and whether such changes might depend on the high endogenous level of DJ-1. We first suppressed endogenous DJ-1 by siRNA and compared these cells with the non-specific siRNA control. Differential miRNA expression analysis showed that DJ-1 depletion produced essentially no detectable alteration in the miRNA profile relative to the untreated cells (**Fig. 1A**; Supplementary **Fig. S2**). Although the siRNA manipulation itself produced measurable transcriptional effects, these effects were not accompanied by a corresponding change in the miRNA repertoire, consistent with a transient response to siRNA introduction and potential activation of innate immune pathways (Oh & Mouradian, 2018).

**Figure 1.**
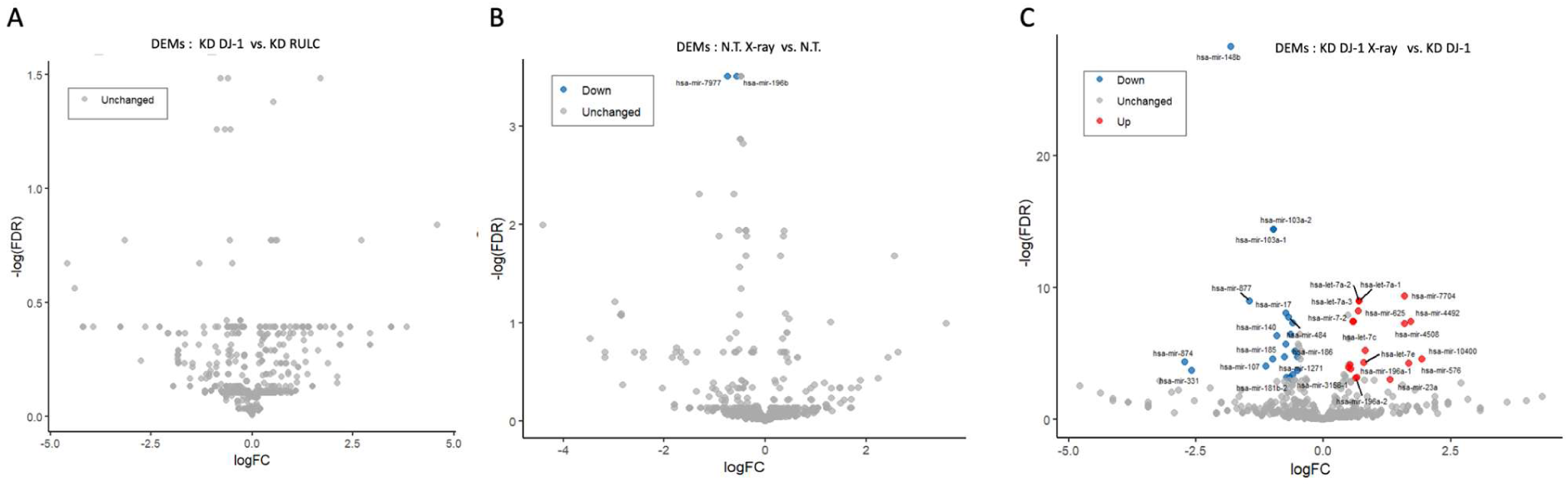
Volcano plots of miRNA expression changes across cellular conditions. Volcano plots show log₂ fold change (logFC) versus log₂(FDR). Upregulated miRNAs are shown in red, downregulated miRNAs in blue, and unchanged miRNAs in grey. Note that axis scales differ between panels. **(A)** DJ-1 knockdown (KD) versus non-specific siRNA control (RULC). **(B)** Naïve cells versus cells exposed to X-ray irradiation. **(C)** miRNA changes in DJ-1 KD cells following X-ray irradiation relative to non-irradiated DJ-1 KD cells. Selected miRNAs are labeled.

We next examined the effect of X-ray irradiation on the miRNA profile of naïve cells retaining endogenous DJ-1. Despite robust activation of the DDR following DNA damage (Zohar et al, 2025), irradiation produced remarkably little change in miRNA expression (**Fig. 1B**). Under these conditions, cell viability remained >95% at 6 h and >90% at 24 h after irradiation, indicating that the cells largely tolerated the treatment. Among 542 uniquely identified mature miRNAs, only miR-7977 and miR-196b-5p showed significant changes, both being modestly downregulated (FDR = 0.03). miR-7977 has previously been implicated in regulation of SIRT3 and ROS levels (Wlodarski et al, 2020). although the modest change observed here is unlikely to account for the overall cellular response.

Importantly, the absence of a substantial miRNA response was also observed following DJ-1 depletion. Despite an approximately eightfold reduction in DJ-1 expression achieved by siRNA (Zohar & Linial, 2024), the mature miRNA profile remained largely unchanged following X-ray exposure (Supplementary **Table S1**). Thus, at this early post-irradiation time point, neither basal DJ-1 expression nor its depletion resulted in a major change in the steady-state mature miRNA repertoire. These findings established a robust baseline where the mature-miRNA repertoire is remarkably stable in both naïve and DJ-1-depleted cells. This observation prompted us to examine whether DJ-1 instead influences the capacity to remodel the miRNA system by genotoxic stress by X-ray and monitored the cells 6 h post irradiation.

### Suppression of DJ-1 by siRNA increases sensitivity to X-ray irradiation and modestly alters miRNAs

To assess the impact of DJ-1 depletion on the miRNA response to ionizing radiation, we profiled miRNA expression in DJ-1 knockdown (KD) cells 6 h following X-ray irradiation. The complete list of differentially expressed miRNAs (DEMs) is provided in Supplementary **Table S2**. Among the 542 uniquely detected miRNAs, only a limited subset met the criteria for differential expression. As shown in **Fig. 1C**, 14 miRNAs were upregulated and 20 were downregulated. Among the upregulated miRNAs, highly expressed species such as miR-7-5p (representing >1% of total cellular miRNAs) and let-7a-5p showed consistent but modest increases of 1.5-fold and 1.6-fold, respectively. These changes may reflect an adaptive response to radiation-induced stress, potentially through coordination of the DNA damage response (DDR) or modulation of pro-survival signaling pathways.

Several moderately downregulated miRNAs, including miR-151a, miR-103a, and miR-148b, maintained intermediate to high basal expression levels. Additional downregulated miRNAs with lower abundance, such as miR-331 and miR-874, further indicate that irradiation of DJ-1-depleted cells affects a relatively restricted subset of the miRNA repertoire. Overall, only 6.3% of detected miRNAs were differentially expressed, and only 2% met the more stringent threshold of |log₂(FC)| ≥ 1.0. Moreover, the magnitude of change among significant DEMs was generally low to modest. These findings are consistent with a limited adaptive response to irradiation rather than a broad miRNA reprogramming event at 6 h post-irradiation.

### Network analysis of DEM targets following irradiation in DJ-1-depleted cells reveals enrichment of cancer-related pathways

To explore the functional implications of the DEMs in DJ-1-depleted cells following irradiation, we analyzed shared miRNA–target networks and performed KEGG pathway enrichment analysis. **Fig. 2A** presents a network of the 14 upregulated miRNAs and their validated targets, highlighting genes targeted by multiple miRNAs. Several highly connected target genes (pink nodes) and represent candidate regulatory hubs subject to coordinated miRNA regulation. **Fig. 2B** shows the connectivity network of the 20 downregulated miRNAs and their validated targets. In contrast to the upregulated miRNA network, the downregulated miRNA–target network was relatively sparse, with limited connectivity and no prominent regulatory hub.

**Figure 2.**
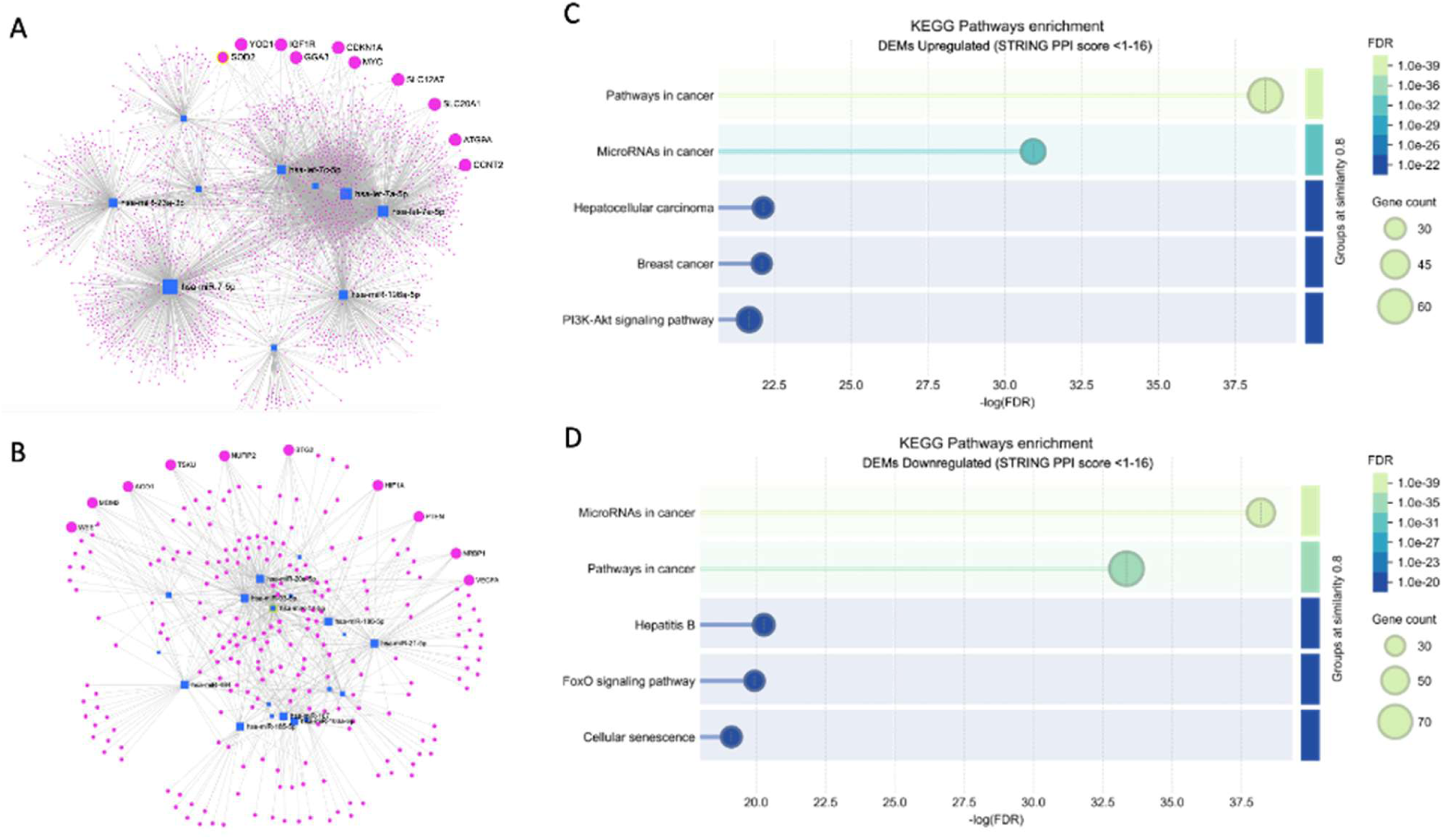
Network and pathway analysis of miRNA changes in DJ-1 knockdown cells after irradiation. **(A)** Network of upregulated DEMs and their validated targets (miRTarBase v9.0; miRNet 2.0). Nodes represent miRNAs and target genes; pink nodes indicate highly connected targets regulated by multiple miRNAs. **(B)** Network of downregulated DEMs and their validated targets, showing sparse connectivity without a dominant miRNA hub. **(C)** STRING protein–protein interaction (PPI) network of validated targets of upregulated DEMs showing significant connectivity (PPI < 1 × 10⁻¹⁶) and enrichment for cancer-associated pathways. **(D)** KEGG pathway enrichment analysis of validated targets highlighting cancer-related signaling and miRNA-regulated pathways.

Experimentally validated targets from miRTarBase v9.0 were further analyzed using STRING to assess functional enrichment. As shown in **Figs. 2C** and **2D**, the resulting networks showed significant enrichment for cancer-related signaling pathways, including pathways associated with miRNA-mediated regulation in cancer. Thus, although the number of affected miRNAs was limited (34 DEMs in total) and their fold changes were generally modest, their predicted targeting of central regulatory genes suggests that the response of DJ-1-depleted cells to irradiation may nevertheless influence regulatory pathways commonly dysregulated in cancer.

### X-ray irradiation promotes global alterations in miRNA expression levels

We next challenged the cellular response to X-ray irradiation under conditions of DJ-1 overexpression (OX). DJ-1 was overexpressed using a DJ-1 expression plasmid (using empty-plasmid control as reference; Supplementary **Table S3**). We then focused on miRNAs that were significantly altered following X-ray treatment (Supplementary **Table S4**). **Fig. 3A** shows that the fraction of differentially expressed miRNAs (DEMs) in the DJ-1 OX background was substantially higher than that observed following X-ray treatment of cells expressing endogenous DJ-1 (37.2% DEMs). Thus, elevated DJ-1 markedly increased the breadth of the miRNA response to irradiation. This effect was further evident when considering the distribution of total miRNA reads **(Fig. 3B)**. Of the 716 identified miRNAs, more than half of all reads were associated with downregulated miRNAs following irradiation, whereas only ∼11% of reads were assigned to miRNAs classified as unchanged (see Methods).

**Figure 3.**
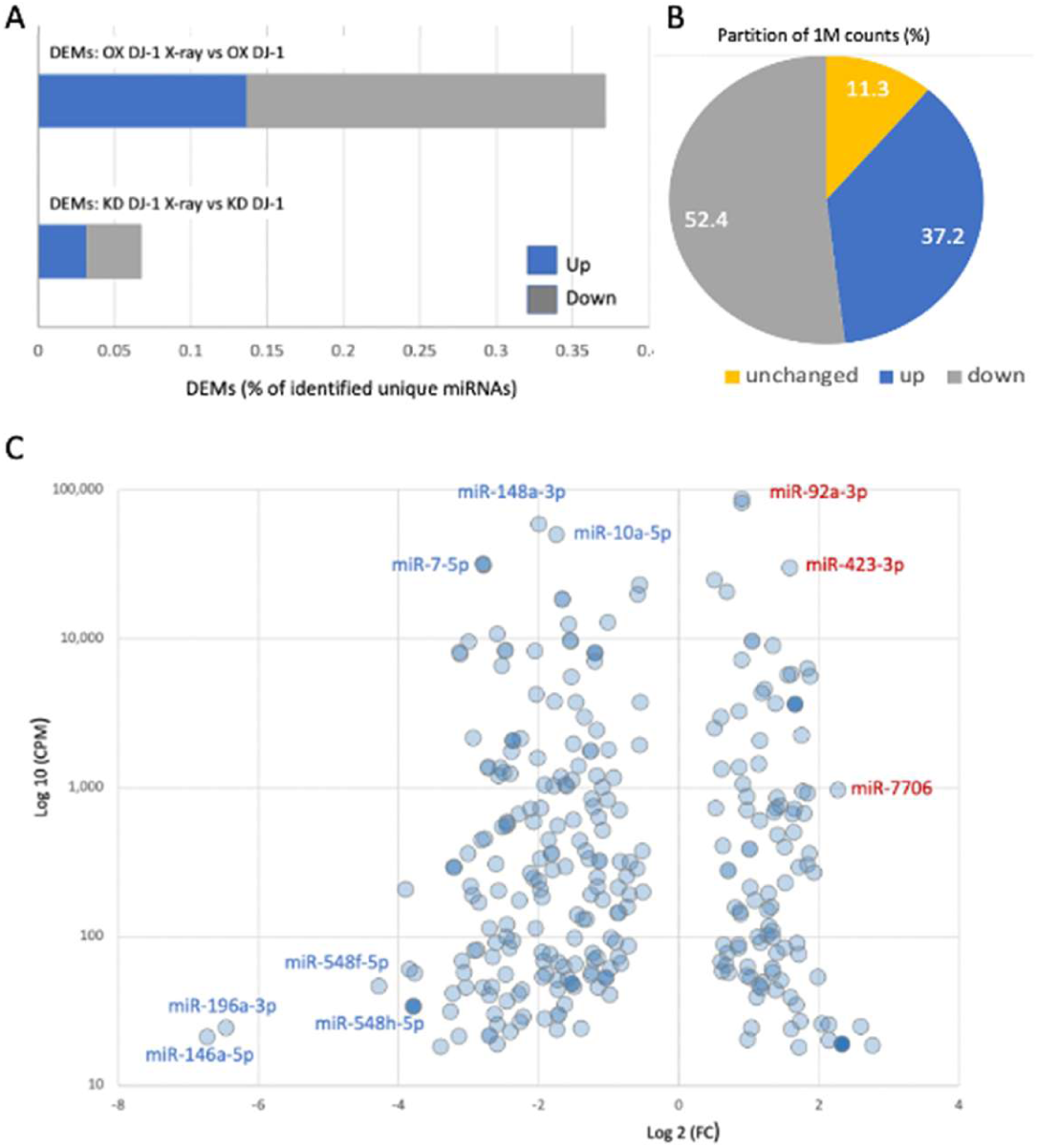
X-ray–induced miRNA expression changes in DJ-1-overexpressing cells. **(A)** Proportion of differentially expressed miRNAs (DEMs; upregulated and downregulated) following X-ray irradiation in DJ-1-overexpressing cells (DJ-1 OX X-ray) relative to DJ-1 OX cells. Bars represent the percentage of DEMs relative to the total number of detected unique miRNAs using a threshold of >18 nCPM. **(B)** Distribution of 1M reads across the 716 identified miRNAs (Supplementary **Table S4**), according to the expression trends following X-ray irradiation in DJ-1 OX cells (upregulated, downregulated, or unchanged). **(C)** Scatter plot of miRNA expression changes in DJ-1 OX cells following X-ray irradiation relative to DJ-1 OX cells, showing log₂ fold change (FC) versus log₁₀ normalized expression (CPM). Selected deregulated miRNAs are labeled.

The comparison of the number of affected miRNAs with their contribution to the total read pool indicates that the response was driven primarily by abundance miRNA species. Many of the 716 detected miRNAs were classified as unchanged (371 miRNAs), largely because they failed to meet the predefined minimum expression threshold (≥18 nCPM) and/or statistical significance criterion (FDR <0.05). In contrast, several highly abundant miRNAs showed pronounced changes following irradiation. These included miR-146a, miR-106a, and miR-7, which were strongly downregulated. These miRNAs have previously been implicated in inflammatory signaling, oxidative stress responses, and cell-cycle regulation. In contrast, miR-92a-3p, miR-423-3p, and miR-7706 were among the most strongly upregulated miRNAs following irradiation, suggesting activation of distinct adaptive regulatory programs (**Fig. 3C**).

Several members of the miR-548 family, including miR-548f-5p and miR-548h-5p, were strongly downregulated following irradiation, with reductions of approximately 13–14-fold. Even more pronounced suppression was observed for miR-146a-5p and miR-196a-3p, which decreased by approximately 88–100-fold. These miRNAs have been linked to stress- and immune-related regulatory pathways, suggesting that their marked suppression may contribute to the remodeling of stress-responsive regulatory programs following X-ray exposure.

Together, these data indicate that DJ-1 overexpression markedly increases the dynamic range of the miRNA response to X-ray irradiation, resulting in extensive remodeling of the miRNA landscape. This response contrasts sharply with the limited miRNA changes observed following DJ-1 KD and suggests that DJ-1 levels influence the capacity of cells to reprogram their miRNA repertoire in response to genotoxic stress. A complete list of DEMs identified following X-ray irradiation in the DJ-1 OX and empty-plasmid control settings is provided in Supplementary **Table S4.**

Similar to the general distribution of miRNA abundance across cells, a small number of miRNAs account for a large proportion of total miRNA reads. In our dataset, 64% of all reads were contributed by 18 most abundant miRNAs, with miR-92a-3p alone accounting for 16.7% of the total cellular miRNA reads. Importantly, differential expression of highly abundant miRNAs (defined here as ≥1% of total reads per miRNA) is likely to have a disproportionate effect on the overall composition of the cellular miRNA pool, indirectly reshaping the relative abundance of the remaining miRNAs (Mahlab-Aviv et al, 2019). In this context, miR-17-5p, miR-26a-5p, miR-7-5p, miR-10a-5p, and miR-148a-3p together accounted for more than 24% of the total miRNA reads and were strongly downregulated following X-ray irradiation. Suppression of these highly abundant miRNAs may therefore have a coordinated impact on cellular stress responses. Under steady-state conditions, these miRNAs regulate pathways involved in cell-cycle progression, PI3K–AKT and EGFR signaling, and metabolic homeostasis. We conclude that DJ-1 overexpression enables broad remodeling of the mature miRNA landscape following DNA damage, whereas DJ-1 depletion is associated with a markedly blunted and minimal miRNA response.

### The role of miRNAs as mediators for genotoxic gene expression rebalancing

In our previous study (Zohar et al, 2025) we observed that upon manipulation of DJ-1 under a genotoxic stress, cells activate quite a substantial change in ncRNAs. We asked whether the observed changes in miRNAs (this study) across all cellular conditions can be explained at a system level. Namely, whether the data support a contribution of miRNAs in reshaping cell state. To this end, we used a global view of the different experimental settings using principal component analysis (PCA). PCA results derived from analyzing miRNA across several cellular settings of DJ-1 OX and DJ-1 KD conditions in responses to X-ray stress are shown (Supplementary **Fig. S3).** Under DJ-1 knockdown, miRNA variance is very modest (PC1 18%) suggesting that miRNA dynamics is ineffective in propagating the stress-induced gene expression changes. In contrast, for DJ-1 OX setting accounts for a substantial fraction of the explained variance. The PC1 for miRNAs for the setting of OX DJ-1 explains 55% of the variance. This pattern is consistent with a scenario in which miRNA changes represent a dominant layer of regulation following the genotoxic stress response, where miRNAs seem to contribute mostly to the DJ-1 OX status.

### Gene targets collectively regulated by upregulated DEMs are involved in cellular metabolism and homeostasis

We next examined the potential collective effects of the upregulated DEMs by identifying their most likely target genes. Using the miEAA tool (see Methods), we identified 377 enriched target genes, ranked according to their enrichment scores (Supplementary **Table S5**). The top 26 enriched genes are shown in **Fig. 4A**. We further selected the top miRNAs (q-value <1.0 × 10⁻⁵) for STRING functional protein–protein interaction (PPI) network analysis, yielding a network of 43 genes **(Fig. 4B)**. The resulting network showed significant enrichment for RNA-binding proteins and cellular metabolic processes. It was dominated by ribosomal proteins (e.g., RPL8, RPL26, RPS2, and RPL3), translation-related factors (EEF1A1, EIF3I, and GCN1), and proteins involved in mitochondrial metabolism and respiration (MT-ND1, MT-ND6, and MT-CO1). These findings are consistent with coordinated miRNA-mediated regulation of translational activity, mitochondrial function, and cellular metabolic homeostasis following genotoxic stress.

**Figure 4.**
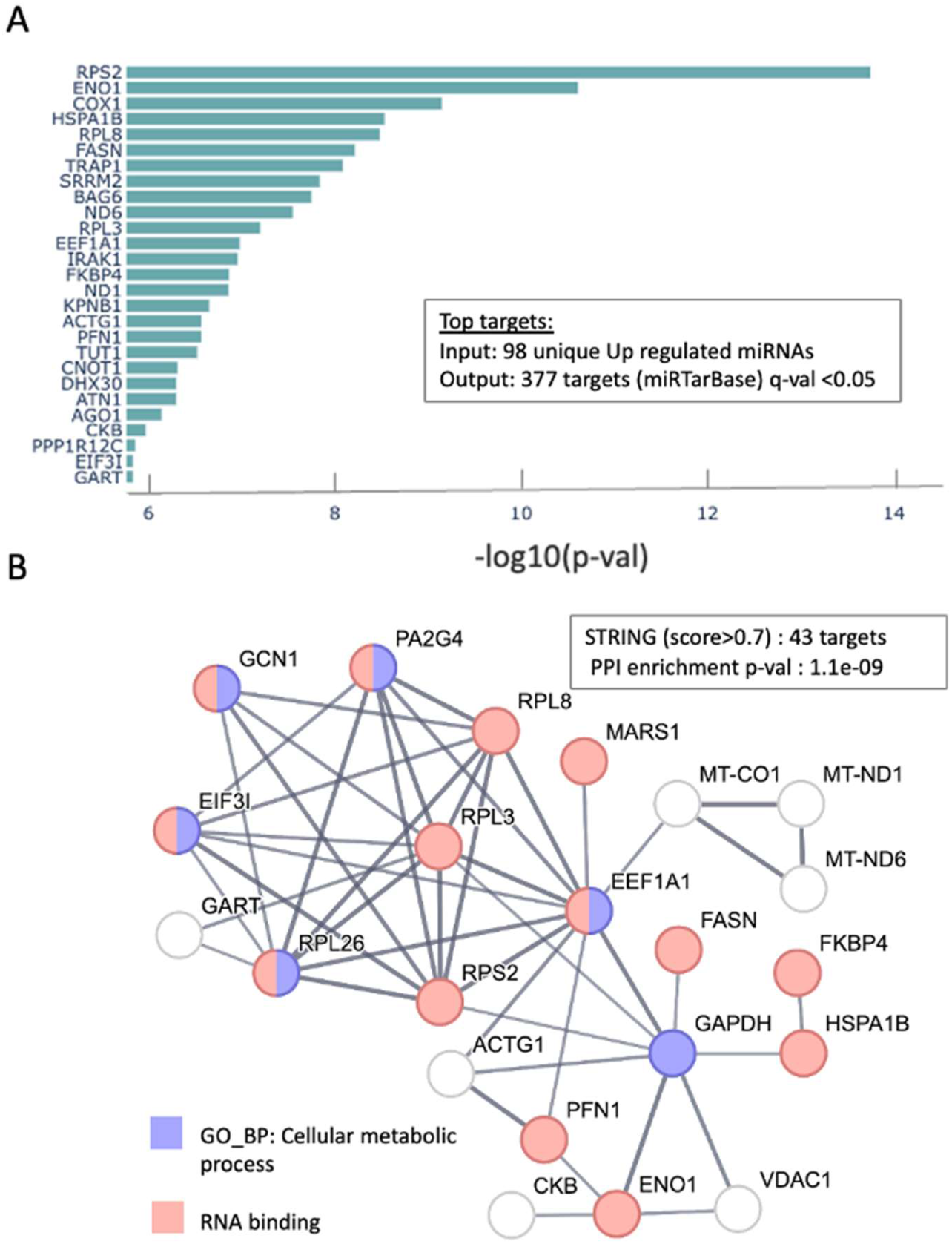
Top targets of miRNAs upregulated by genotoxic stress in a DJ-1 OX background. **(A)** DEMs upregulated upon X-ray (Supplementary **Table S4**) irradiation in DJ-1–overexpressing cells were analyzed for experimentally supported targets using miRTarBase. From 98 unique upregulated miRNAs, 377 significant target genes were identified (q-value <0.05, Supplementary **Table S5**). Enrichment ranking of the most significant target genes based on –log10(p-value). **(B)** Protein-protein interaction (PPI) network of the top targets (43, q-val <1.0e-05) visualized using STRING (interaction score > 0.7; PPI enrichment p-value 1.1e-09). Nodes are colored by functional annotation.

The genes involved in energy homeostasis (COX1, ND1, ND6, ENO1, VDAC1, and GAPDH) and RNA processing and miRNA regulation (SRRM2, AGO1, CNOT1, TUT1, and RBM10) further support a role for miRNAs in maintaining cellular metabolic and regulatory homeostasis. In addition, stress-responsive chaperones (HSPA1B, TRAP1, and BAG6), cytoskeletal and trafficking regulators (ACTG1, PFN1, KPNB1, and VPS18), and signaling mediators (IRAK1 and HIPK2) were highlighted, supporting a model in which radiation-induced miRNA changes contribute to cellular stress-adaptation pathways. Overall, this pattern of miRNA-mediated regulation is consistent with reported radiation responses involving coordinated changes in metabolism, translation, and RNA processing that promote cellular adaptation to DNA damage (Podralska et al, 2020).

### Gene targets collectively regulated by downregulated DEMs are enriched in RNA metabolism

The majority of miRNAs were downregulated following X-ray irradiation **(Fig. 5).** To identify the potential targets and pathways collectively affected by these miRNAs, we analyzed the 168 uniquely identified downregulated miRNAs, which yielded 238 target genes (Supplementary **Table S5).** To enable a direct comparison with the upregulated miRNA analysis **(Fig. 4),** we matched the input size for the STRING network analysis **(Fig. 5A)** and for KEGG and WikiPathways enrichment analyses **(Figs. 5B, 5C).** Several findings emerged from this analysis. First, the most significantly enriched KEGG pathway was miRNA in cancer, and DNA damage response (DDR) was the most significantly enriched term among the WikiPathways. Second, several core regulators of miRNA biogenesis and RNA metabolism emerged among the most extensively targeted genes. DICER1 was targeted by 30 of the input miRNAs (q-value = 7.90 × 10⁻¹⁴), and AGO1 was targeted by 34 miRNAs (q-value = 4.21 × 10⁻¹²), indicating broad coordinated regulation of key components of the miRNA-processing and effector machinery.

**Figure 5.**
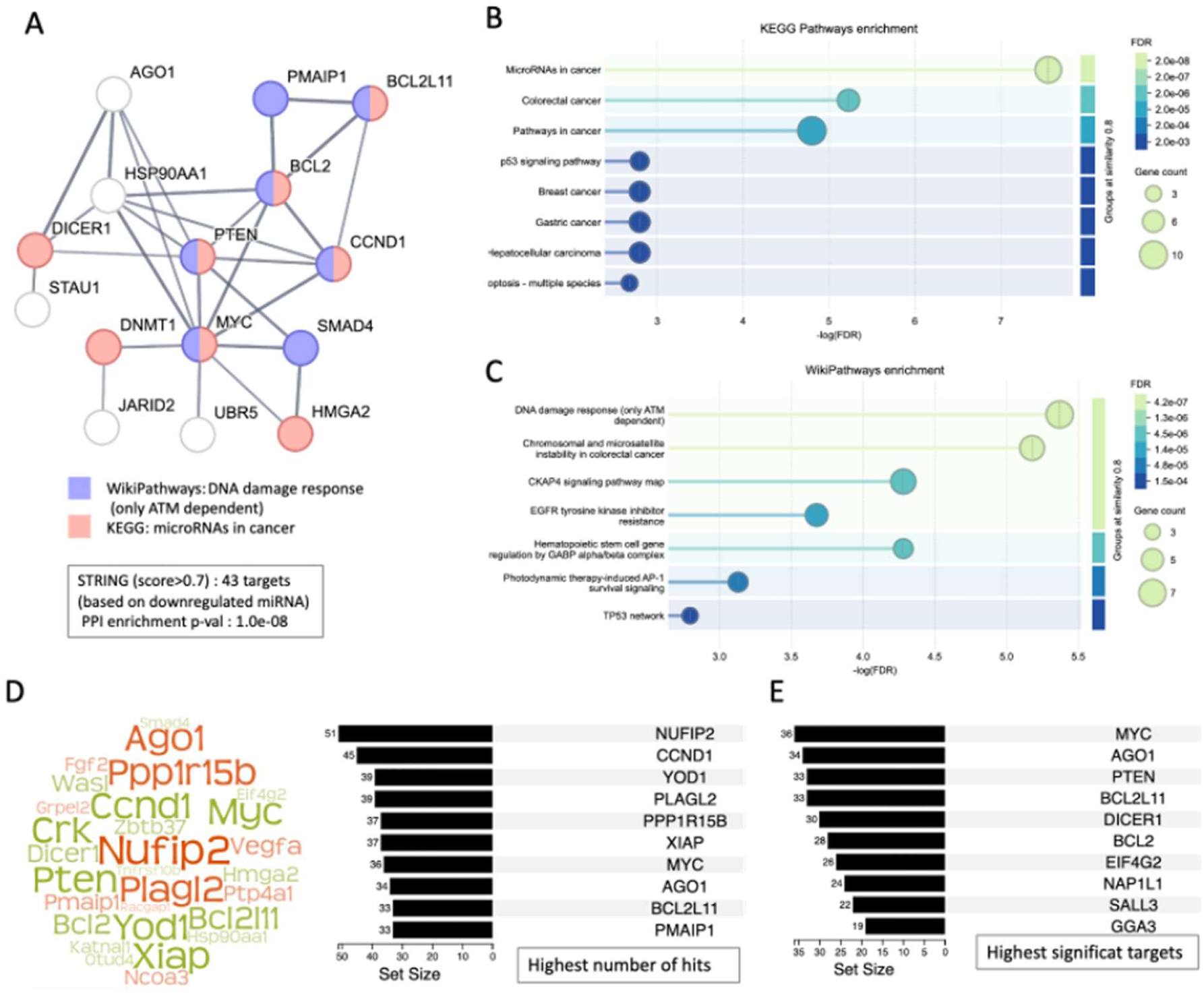
Network and pathway enrichment analysis of targets of miRNAs downregulated by X-ray irradiation in DJ-1 overexpressing cells. **(A)** Top target genes identified by miEAA (minimum FDR <1 × 10⁻⁴) were selected for STRING PPI and functional enrichment analyses. The resulting PPI network (PPI enrichment p-value = 1 × 10⁻⁸) highlights functional associations related to the DNA damage response (DDR) and miRNA-mediated regulation in cancer. **(B)** Pathway enrichment analysis based on KEGG. **(C)** Pathway enrichment analysis based on WikiPathways. Node colors and sizes indicate coverage and enrichment significance by −log₁₀(FDR). **(D)** Word cloud of the top target genes regulated by the largest number of downregulated DEMs (left); larger font size indicates regulation by a greater number of miRNAs. The top 10 targets are listed on the right. **(E)** Top 10 target genes ranked by statistical significance (q-value), together with the number of downregulated miRNAs predicted to regulate each gene.

A particularly prominent target was NUFIP2 (nuclear fragile X mental retardation protein-interacting protein 2), which was targeted by 51 of the 168 input miRNAs. NUFIP2 is an RNA-binding adaptor involved in RNA metabolism, translational control, and cellular stress responses, including functions associated with ribonucleoprotein (RNP) granules and miRNA-mediated regulation (Bish et al, 2015). YOD1 was also among the most extensively targeted genes (39 miRNAs; q-value = 4.89 × 10⁻⁸). YOD1 encodes a regulatory deubiquitinase involved in the extraction and processing of misfolded proteins from the endoplasmic reticulum (ER). Its activity is particularly relevant when translation or protein folding is perturbed, helping cells cope with increased loads of damaged or misfolded proteins and maintain cellular homeostasis (Tanji et al, 2018).

**Fig. 5D** shows a word-cloud of most likely targets that match the input miRNA set, ranked according to the number of miRNA binding site hits, listing the top 10 targets highly regulated by diverse number of miRNAs according to miRTarBase experimental results. **Fig. 5E** lists the genes that are statistically significant with 10 top targets. Note that MYC, AGO1, BCL2L11 were predicted in both top target lists (**Figs. 5D** and **5E**). Upregulation of MYC points to activation of mitogenic signaling and transcriptional amplification, promoting ribosome biogenesis, metabolic activity, and cell cycle progression.

Of particular interest is AGO1 (Argonaute 1) that emerged as a prominent target of the radiation-responsive miRNA network. AGO1 was among the most significantly enriched targets of the downregulated miRNAs following X-ray irradiation in DJ-1-overexpressing cells, with multiple miRNAs collectively predicted to regulate its expression. This observation identifies AGO1 as a potential regulatory hub within the radiation-responsive miRNA network (**Figs. 5D**, **5E**). The observed changes may reflect a shift in post-transcriptional regulation toward buffering and fine-tuning of gene expression. This interpretation is consistent with the enrichment in metabolic rebalancing **(Figs. 5B, 5C).**

MiEAA 3.0 analysis further revealed a highly coordinated targeting pattern among the downregulated DEMs in the DJ-1 OX X-ray versus DJ-1 OX comparison. Among the downregulated DEMs (259 miRNAs targeting 222 unique genes), each target was regulated by an average of 33 miRNAs (Supplementary **Fig. S4**), suggesting preferential and potentially cooperative regulation of these transcripts under genotoxic stress. Cross-referencing these predictions with experimental AGO-CLIP data from ENCORI (Zhou et al, 2026) provided a similar high interactions per gene (on average, 29 experimental validated). Notably, MYC, NAPIL1, and PPP1R15B each is supported by >60 experimental AGO-CLIP interactions.

In contrast, the upregulated DEM set (149 miRNAs targeting 126 unique genes) showed substantially less convergence, with an average of 11 miRNAs per target gene (14 based on experimental AGO-CLIP interactions per gene; Supplementary **Fig. S4).** Together, these findings indicate that X-ray irradiation in the context of elevated DJ-1 produces a pronounced rewiring of post-transcriptional regulation, with the strongest signal arising from the coordinated downregulation of miRNAs. The emergence of highly connected target genes involved in stress responses, mitochondrial regulation, and proteostasis further suggests that miRNA-mediated post-transcriptional control contributes to shaping the cellular response to DJ-1-associated oxidative and genotoxic stress. Supplementary **Table S5** lists the experimentally supported miRTarBase targets of the affected miRNAs in the DJ-1 KD X-ray versus DJ-1 KD comparison, separated into upregulated and downregulated miRNAs.

### DJ-1 determines the extent of miRNA remodeling following genotoxic stress

To reach the mechanistic underlying explanation for the substantial difference in the nature and the amounts of DEMs between DJ-1-depleted and DJ-1-overexpressing cells following X-ray exposure (e.g., **Fig. 3A**), we the two experimental settings with DJ-1 OX and KD were compared in view of the miRNA machinery. We examined the mRNA expression levels of 31 transcripts involved in miRNA biogenesis, maturation, RISC assembly, RNA remodeling, and mRNA decay following X-ray irradiation. These gene products were assigned to functional modules encompassing (i) nuclear pri-miRNA processing, (ii) nuclear export of pre-miRNAs, (iii) Dicer-dependent maturation, (iv) miRNA stability and maturation regulators (v) Argonaute proteins, (vi) RISC effector proteins, (vii) RNA helicases and remodeling factors, (viii) P-body components, and (ix) RNA-binding stress-response proteins.

The response of the miRNA machinery to X-ray irradiation was markedly different following DJ-1 OX and KD (Supplementary **Table S6**). In DJ-1-depleted cells, irradiation produced a coordinated suppression of multiple components of canonical miRNA biogenesis. DICER1, the central RNase III enzyme responsible for cleavage of pre-miRNAs, was reduced by approximately 60% (p=0.001). Its cofactor PRKRA (PACT) was reduced by approximately 44% (p=0.001). Two terminal uridylyl transferases involved in miRNA maturation and turnover, TUT4 and TUT7, were reduced by 60% each (p<0.001 and p=0.003, respectively). We observed that additional components acting at earlier stages of miRNA processing were also suppressed, including DDX17 (∼28% reduction, p=0.007) and SMAD2 (∼30% reduction, p=0.004). LIN28B was reduced by 50% (p=0.012), further indicating broad suppression of miRNA maturation pathways (**Table 1**).

**Table 1.**
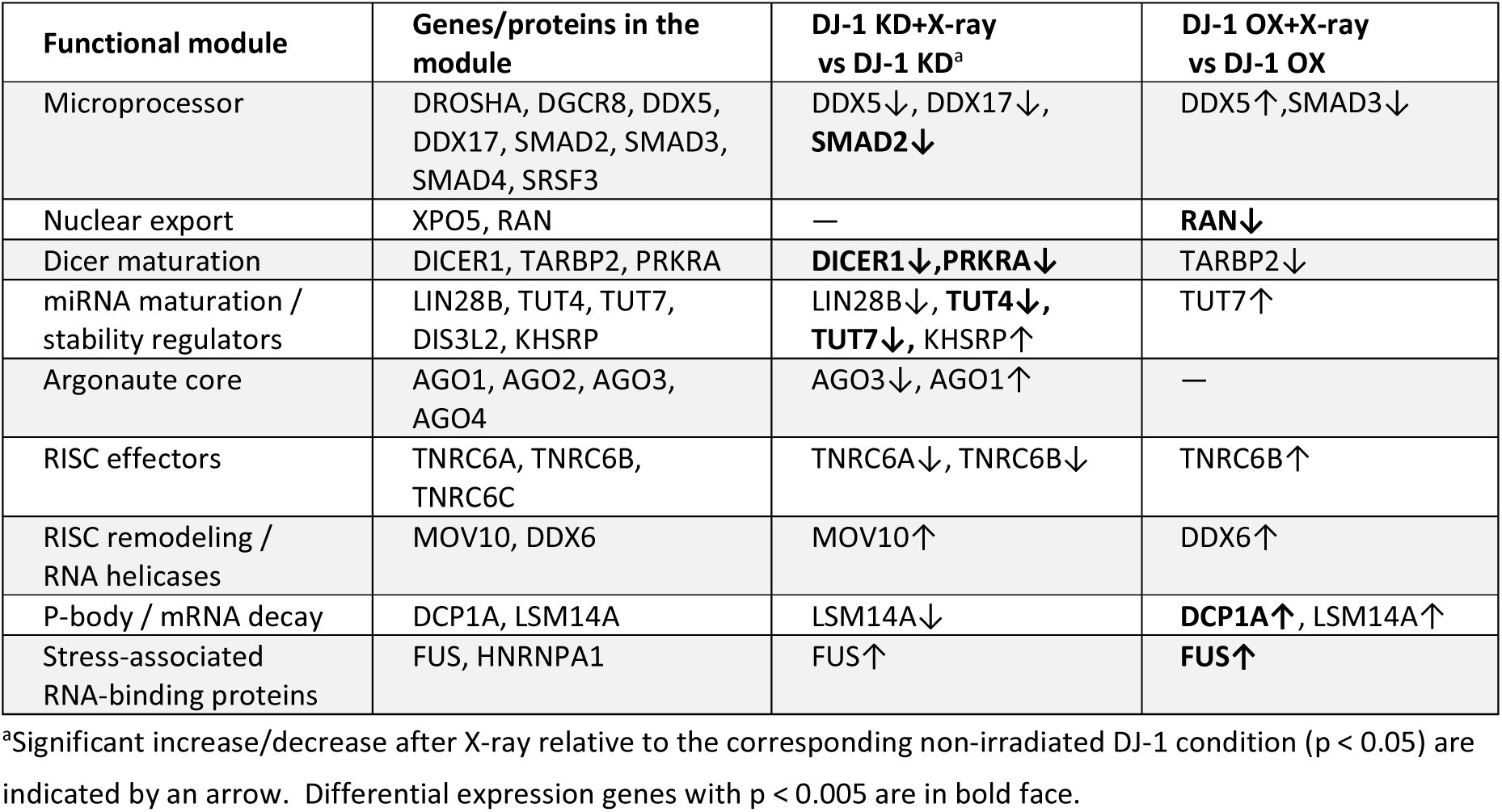
miRNA regulatory machinery following genotoxic stress in DJ-1-deficient and DJ-1-overexpressing cells

| Functional module | Genes/proteins in the module | DJ-1 KD+X-ray vs DJ-1 KD<sup>a</sup> | DJ-1 OX+X-ray vs DJ-1 OX |
| --- | --- | --- | --- |
| Microprocessor | DROSHA, DGCR8, DDX5, DDX17, SMAD2, SMAD3, SMAD4, SRSF3 | DDX5↓, DDX17↓, **SMAD2↓** | DDX5↑, SMAD3↓ |
| Nuclear export | XPO5, RAN | — | **RAN↓** |
| Dicer maturation | DICER1, TARBP2, PRKRA | **DICER1↓, PRKRA↓** | TARBP2↓ |
| miRNA maturation / stability regulators | LIN28B, TUT4, TUT7, DIS3L2, KHSRP | LIN28B↓, **TUT4↓**, **TUT7↓**, KHSRP↑ | TUT7↑ |
| Argonaute core | AGO1, AGO2, AGO3, AGO4 | AGO3↓, AGO1↑ | — |
| RISC effectors | TNRC6A, TNRC6B, TNRC6C | TNRC6A↓, TNRC6B↓ | TNRC6B↑ |
| RISC remodeling / RNA helicases | MOV10, DDX6 | MOV10↑ | DDX6↑ |
| P-body / mRNA decay | DCP1A, LSM14A | LSM14A↓ | **DCP1A↑**, LSM14A↑ |
| Stress-associated RNA-binding proteins | FUS, HNRNPA1 | FUS↑ | **FUS↑** |

<sup>a</sup>Significant increase/decrease after X-ray relative to the corresponding non-irradiated DJ-1 condition ( $p < 0.05$ ) are indicated by an arrow. Differential expression genes with $p < 0.005$ are in bold face.

The suppression extended beyond miRNA biogenesis to components of the effector machinery. AGO3 was reduced by ∼52% (p=0.011), while TNRC6A and TNRC6B, which mediate recruitment of downstream mRNA-silencing and decay machinery to AGO-containing complexes, were reduced by approximately 35% and 39%, respectively (p=0.049 and p=0.036). LSM14A, a component of RNA granules and P-body-associated mRNA decay machinery, was also reduced by approximately 40% (p=0.020). In contrast, AGO1 but not the other AGO genes, showed a modest but significant increase (p=0.012), whereas no genes of the nuclear export system were unaffected (XPO5, RAN). We concluded that among the set of genes linked to miRNA processing, trafficking, and stability, most genes were downregulated following X-ray including specific components of canonical miRNA maturation and the RISC effector pathways **(Table 1).**

A small group of RNA-binding and RNA-remodeling proteins showed the opposite behavior. MOV10 exhibited the strongest induction in the DJ-1-depleted cells following irradiation, corresponding to an approximately 1.9-fold increase with modest statistics (p=0.027) similar to KHSRP and FUS (p-values 0.037 and 0.023, respectively; Supplementary **Table S6**). These proteins have broader roles in stress-associated RNA regulation.

The pattern observed following DJ-1 OX was strikingly different. In contrast to the broad suppression of miRNA-processing components observed following DJ-1 depletion, the components of the canonical miRNA processing pathway were maintained intact after X-ray exposure **(Table 1)**. RAN, which provides the GTPase-dependent transport function required for nuclear export of pre-miRNAs, was significantly reduced (p=0.001), while TARBP2, a major Dicer-associated cofactor, was reduced by approximately 30% (p=0.006). We found that in DJ-1 OX setting following X-ray, the upstream miRNA-processing machinery is preserved, while gene products acting downstream in RNA silencing and RNA decay are affected. Specifically, DCP1A and FUS were slightly but significantly increased (10-20%) and these changes were significant (p=0.003 and p=0.004, respectively) (Supplementary **Table S6**). These proteins participate in RISC-associated silencing, P-body organization, RNA remodeling, decapping, and mRNA turnover. In the context of your DJ-1 and X-ray experiment, FUS increases significantly after X-ray in both DJ-1 KD and DJ-1 OX cells. FUS is an RNA-binding protein with an important role in RNA processing, RNA stability, and stress responses (Shelkovnikova et al, 2014). We conclude that following DNA damage, DJ-1 OX preferentially affected the downstream execution and remodeling of RNA silencing rather than affecting the miRNA biogenesis per se.

### Cellular composition of mature and precursor miRNAs is DJ-1 dependent

To determine whether the transcriptional changes in miRNA-processing pathways were reflected in the composition and organization of the miRNA pool, we next examined miRNA precursors and mature miRNAs. This analysis revealed marked differences in the breadth and composition of the miRNA response between DJ-1-depleted and DJ-1-overexpressing cells. Although the two conditions shared substantial overlap in precursor identity, the DJ-1 OX setting displayed a broader precursor repertoire (Supplementary **Table S7**). We identified 215 precursor miRNAs in DJ-1-depleted cells, of which 94% were also detected in DJ-1-overexpressing cells, which contained 289 unique precursors in total, including 86 unique precursors detected only in the DJ-1 OX setting. In contrast, only 12 precursors were unique to the DJ-1 KD condition (Supplementary **Table S8**). These findings indicate that DJ-1 overexpression does not establish a fundamentally distinct precursor repertoire, but rather is associated with greater breadth and diversity of the precursor miRNA pool.

To determine whether the extensive downregulation of mature miRNAs observed following X-ray irradiation in the DJ-1 OX setting reflected altered miRNA transcription, processing, or stability, we compared miRNA expression at two regulatory levels: precursor transcripts quantified from total RNA-seq (>200 nt) and mature miRNAs quantified by sncRNA-seq. The 292 miRNA genes identified following X-ray irradiation in the DJ-1 OX setting accounted for approximately 82% of the total reads assigned to other noncoding RNA species. Notably, miRNA precursors constituted 71.4% of the statistically significant ncDEGs, indicating a prominent representation of miRNA genes among the radiation-responsive noncoding transcripts.

Precursor expression was analyzed at the gene level (e.g., MIR148A), allowing direct comparison between miRNA gene transcription and the abundance of corresponding mature miRNA products (both miRNA arms). This comparison provides a means to distinguish transcriptional regulation, reflected by concordant precursor: mature changes, from post-transcriptional regulation involving miRNA processing, maturation, or stability. Among the downregulated precursor genes, 140 unique precursor genes were detected, corresponding to 188 unique mature miRNAs (259 mature sequences after sequence compression; **Fig. 6A**). Conversely, only 23 precursor genes were significantly upregulated, whereas 115 unique mature miRNAs were upregulated, corresponding to total 149 mature sequences.

**Figure 6.**
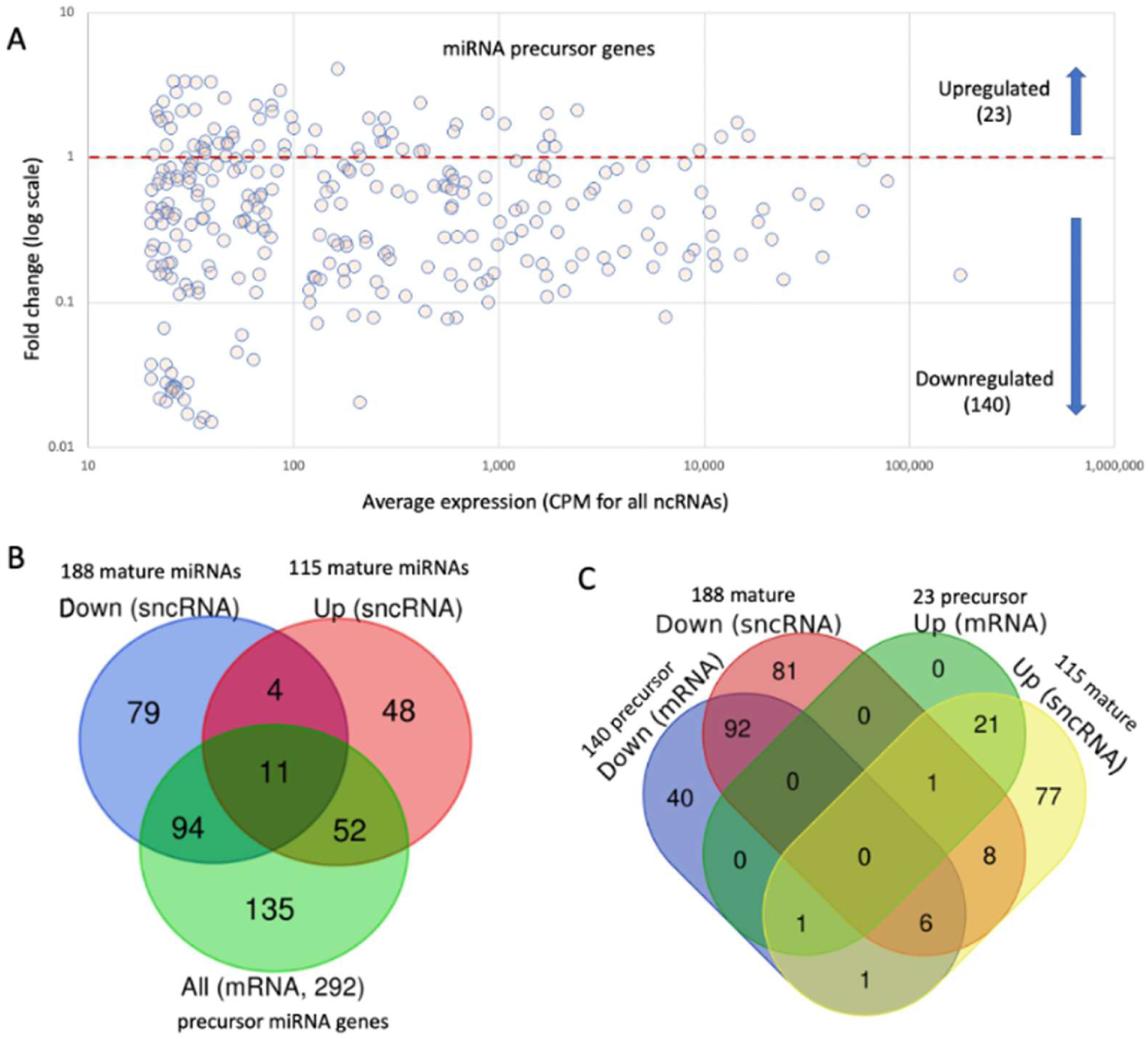
Mapping the expression of miRNA precursor genes. **(A)** An MA plot shows log10(fold change) vs average expression for precursor miRNA genes. The genes are partitioned to up- and downregulated genes. **(B)** Overlap between differentially expressed miRNA precursors (derived from >200 nt total RNA, used for mRNA analysis) and sncRNAs (marked as mature miRNAs), following X-ray exposure in the DJ-1 OX setting. **(C)** Four-way Venn diagram showing the intersection between upregulated (Up) and downregulated (Down) mRNAs and sncRNAs. Numbers denote the count of unique transcripts within each category and their overlaps.

Notably, 91% of the upregulated precursor genes were accompanied by increased levels of mature miRNAs originating from the same genomic loci, consistent with transcriptional induction of these miRNA genes following X-ray irradiation in the DJ-1 OX setting. In contrast, 67% of upregulated mature miRNAs showed no detectable change in their corresponding precursor transcripts, indicating that increased mature miRNA abundance frequently occurred independently of increased precursor production **(Fig. 6B).** This pattern is consistent with DJ-1 that influence the architecture and post-transcriptional dynamics of the miRNA system during stress. In addition, 13% of cases displayed discordant regulation, in which precursor levels decreased while mature miRNA abundance increased. Examples included miR-455, miR-676, miR-532, miR-93, miR-339, and miR-25 **(Fig. 6C)**. Several of these miRNAs are encoded within genomic clusters or represent intronic or arm-flexible miRNAs, and their precursor–mature uncoupling further supports regulation at the level of processing or stability under X-ray-induced, DJ-1-modulated stress.

For downregulated mature miRNAs, 48% showed a corresponding decrease in precursor expression, consistent with transcriptional repression. In contrast, 43% showed no corresponding change in their precursor, indicating that their reduction may instead arise from altered processing, maturation, or stability. Furthermore, 28% of downregulated precursor genes were not accompanied by detectable changes in mature miRNA abundance, providing additional evidence that transcription and mature miRNA abundance can become partially uncoupled following irradiation. For example, miR-191, which accounts for approximately 2.7% of total cellular miRNA abundance, exhibited a twofold decrease in precursor expression together with a twofold increase in mature miRNA levels, providing a clear example of dissociation between transcriptional and post-transcriptional regulation.

A further distinction between the DJ-1 KD and OX settings was evident in the contribution of miRNA precursors to the overall noncoding transcriptome. Following genotoxic stress, precursor-associated reads represented only ∼32% of ncRNA reads in DJ-1-depleted cell sequencing library, compared with **∼**82% in the DJ-1 OX derived sequencing library. Thus, the differences between the two conditions extend beyond changes in individual miRNAs. These findings indicate that DJ-1 influences the cellular architecture of the miRNA system at both the precursor and mature levels. DJ-1 overexpression is associated with a broader precursor repertoire and a pronounced redistribution of the mature miRNA pool following DNA damage, with substantial evidence for regulation beyond transcription alone. In contrast, the relatively stable precursor and mature miRNA landscape in DJ-1-depleted cells is consistent with a markedly more constrained miRNA response over the 6 h post-irradiation interval.

### Radiation-responsive genes show reduced miRNA targeting potential in DJ-1-overexpressing cells

To determine whether the extensive remodeling of the miRNA landscape was reflected in the radiation-responsive coding transcriptome, we examined the distribution of miRNA binding sites across differentially expressed genes (DEGs). In DJ-1-overexpressing cells, the combined set of X-ray-responsive up- and downregulated DEGs contained significantly fewer miRNA binding sites per gene than the background set of coding genes identified by RNA-seq **(Fig. 7A).** The number of binding sites was also lower than that observed for the reference transcription-factor gene set, whereas the difference relative to ribosome-associated genes was not significant **(Fig. 7B).** Thus, despite the extensive remodeling of the mature miRNA landscape following irradiation, the genes undergoing transcriptional changes were not preferentially enriched for miRNA binding sites. Instead, they exhibited a relative depletion of predicted miRNA regulatory sites. A similar analysis for the DJ-1 KD setting resulted in high enrichment of miRNA binding site per DEG when comparted to all identified genes from the RNA-seq of the same conditions (Supplementary **Fig. S5**). Notably, ribosomal genes are almost completely devoid of miRNA binding sites and are no subjected to direct miRNA regulation (**Fig. 7B**). We propose that miRNAs participate in the post-transcriptional adaptation accompanying the translational shutdown in DJ-1 OX, primarily through alternative regulation modes, rather than a direct repression of the bulk ribosomal transcriptome. Potential alternative splicing of transcripts, alteration in 3’-UTR lengths after X-ray, changes in accessibility of by RNA binding proteins are possible molecular mechanisms (Hashemi et al, 2026) that need further inspection.

**Figure 7.**
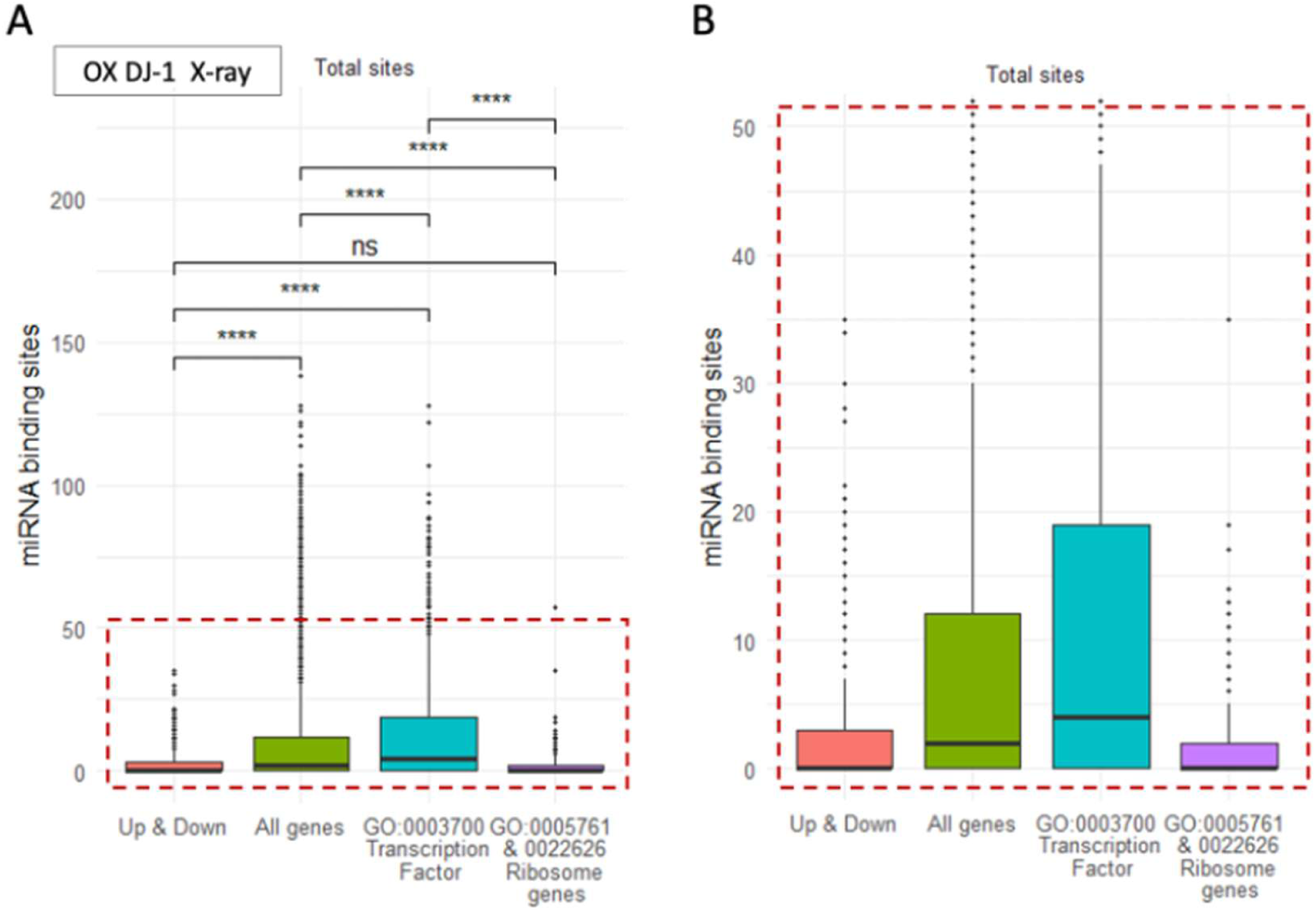
Reduced miRNA binding-site representation among radiation-responsive genes in DJ-1-overexpressing cells. (**A)** Boxplots show the distribution of miRNA binding sites per gene among the combined set of significantly up- and downregulated coding genes following X-ray exposure in DJ-1 OX cells. The distribution is compared with all coding genes identified by RNA-seq and with reference gene sets representing transcription factors (GO:0003700) and ribosome-associated genes (GO:0005761 and GO:0022626). The dashed box highlights the lower range of binding-site distributions. **(B)** Higher resolution for the boxplot results as in the dashed box in A. Statistical comparisons are indicated by **** (p < 1.0 × 10⁻⁴) and ns (p > 0.05).

## Discussion

In this study, we investigated how DJ-1 abundance shapes the cellular response to genotoxic stress, focusing on the contribution of miRNAs to post-transcriptional regulation. Our results support a model in which DJ-1 sets the regulatory capacity and plasticity of the miRNA system following X-ray irradiation. Rather than acting through transcript-specific repression, miRNAs primarily function here as system-level mediators that coordinate translational and metabolic adaptation (Figs. 3 and 4). Our results support the view that rapid changes in relative abundance of large set of miRNAs, especially the abundant ones (Fig. 5) govern cells states and prepare cells and organisms in coping with stress (Biggar & Storey, 2015; Mahlab-Aviv et al, 2021; van Wijk et al, 2022).

The main findings show that the cellular levels of DJ-1 do not determine the basal abundance of miRNAs. Rather, DJ-1 determines the capacity (or plasticity) of the miRNA system to remodel following genotoxic stress. DJ-1 depletion leaves a relatively stable pool of pre-existing mature miRNAs despite evidence for suppression of the processing machinery (Table 1). This observation is consistent with persistence of a relatively stable pool of pre-existing mature miRNAs up to 6 h (Rzeszowska-Wolny et al, 2022), while suggesting reduced capacity for de novo production or remodeling (Mahlab-Aviv et al, 2019). In contrast, DJ-1 overexpression seems to fully preserves the processing system (Table 1) but not miRNA trafficking or the downstream aspects of miRNA stability. Consequently, it enables extensive remodeling of both precursor and mature miRNAs (Fig. 6). These findings extend our previous characterization of the DJ-1–dependent response to X-ray stress, which revealed extensive remodeling of ribosomal, mitochondrial, translational, and lncRNA programs (Zohar et al, 2025). The present study adds miRNA regulation as another layer of this response, with altered miRNA profiles and target enrichment in pathways related to translation, metabolism, and RNA regulation. Together, these findings support a model in which DJ-1 coordinates transcriptional and post-transcriptional remodeling, integrating mRNA, lncRNA, and miRNA changes to adapt translational and metabolic programs to genotoxic and oxidative stress. Cellular adaptation to DNA damage depends not only on transcriptional rewiring but also on dynamic remodeling of miRNA networks. In many cellular contexts, miRNAs were established as agents for keeping cell homeostasis (Olejniczak et al, 2018). Still, the mechanisms coupling stress sensing to miRNA rebalancing remain poorly defined. Our findings identify DJ-1 as a key upstream modulator that licenses miRNA reprogramming specifically under genotoxic stress. The miRNAs within RISC are extremely and expected to act as rapid responder agents. The capacity of miRNAs pools to redistribute among nuclei, cytosol, P-bodies and exosomes, positions them as early regulators, preceding and constraining later mRNA and lncRNA transcriptional programs (van Wijk et al, 2022).

Notably, DJ-1 does not appear to control basal miRNA biogenesis. In naïve HEK293 cells, X-ray irradiation triggered a robust DNA damage response, accompanied by marked transcriptional (mRNA) changes, with minimal miRNA changes (Fig. 1A). Similarly, DJ-1 KD alone did not substantially alter the miRNA landscape which were surprisingly robust, even under X-ray irradiation (Fig. 1B). We show that DJ-1 depleted cells exhibited only very modest miRNA alterations, despite a widespread of thousands of genes and a global mRNA downregulation (Zohar et al, 2025). This transcriptome pattern suggests a fragile cellular state with limited post-transcriptional buffering capacity, rather than coordinated adaptive reprogramming. In striking contrast, DJ-1 overexpression profoundly reshaped the miRNA response to irradiation. While overexpression of DJ-1 alone had negligible effects, irradiation in this context induced widespread miRNA remodeling affecting 37% of detectable miRNAs (nearly 50% with a relaxed expression threshold), and a large majority of all miRNA cell quantities (Fig. 3). PCA analysis demonstrated that miRNA profiles captured a substantial fraction of radiation-induced variance, consistent with miRNAs acting as an early regulatory layer that constrains downstream transcriptional outputs.

We propose that, in a background of excess of DJ-1, irradiation triggers a coordinated wave of miRNAs that modulates translational capacity, resource allocation, and expression noise buffering. The interpretation where DJ-1 overexpression unlocks a large miRNA regulatory response supports the idea that stress signaling must reach a threshold to activate miRNA-mediated buffering. However, it is important to emphasize that DJ-1 depletion and overexpression represent asymmetric perturbations. The pronounced OX phenotype should be interpreted as evidence that increased DJ-1 availability can enable miRNA remodeling, rather than as proof of a dose-dependent relationship.

The mapping of miRNA changes while measuring direct suppression of specific targets in cellular context is limited and prone to false positives and faulty predictions (e.g., (Fridrich et al, 2019; Mockly & Seitz, 2019; Riolo et al, 2020)). We inspected the possible target at a cellular level with the assumption that miRNAs work in cooperation to downregulate targets that have the maximal likelihood to be directly regulated (Balaga et al, 2012). Tools such as miEAA (Aparicio-Puerta et al, 2023) reinforces such systems-level interpretation (discussed in (Blass et al, 2022; Friedman et al, 2014)). To improve reliability in asking who might be the targets that responded to the coordinated changes upon X-ray in OX DJ-1 setting, we only considered experimentally validated findings (miRTarBase, ENCORI). We have identified enriched categories of ribosomal proteins, mitochondrial transcripts, and RNA-binding factors. These gene classes unlikely to be regulated predominantly through direct miRNA binding. Instead, these results suggest indirect control via upstream regulators of translation and metabolism, potentially involving mTOR, MYC, and transcription factors that act as hubs in maintaining homeostasis. Thus, miRNAs appear to modulate metabolic cellular state rather than execute direct transcript-specific repression (Fig. 4).

An additional layer of the response emerged when we examined the potential miRNA regulation of the coding genes that were transcriptionally altered following irradiation in DJ-1-overexpressing cells. Despite the extensive remodeling of the mature miRNA landscape, the X-ray-responsive DEGs showed fewer miRNA binding sites per gene than the background coding transcriptome (Fig. 7). This finding argues against a simple model in which the extensive radiation-induced changes in miRNA abundance directly drive the transcriptional changes observed. Instead, the reduced representation of miRNA binding sites among radiation-responsive genes is consistent with a model in which miRNA remodeling is consistent with a role for miRNAs acting preferentially on transcripts that are not necessarily undergoing strong transcriptional modulation. In this context, the extensive remodeling of the miRNA pool following irradiation may serve to buffer and fine-tune cellular gene expression rather than simply reinforce the transcriptional response. Such a mechanism would be particularly compatible with the DJ-1-overexpressing state, in which the mature miRNA landscape was extensively altered (e.g., RAN, FUS), while the canonical miRNA-processing machinery remained comparatively preserved (Table 1). Together, these observations suggest that DJ-1 may promote a coordinated separation between transcriptional responses to DNA damage and post-transcriptional miRNA regulation, allowing the latter to provide additional flexibility in maintaining cellular homeostasis during genotoxic stress.

A complementary axis emerged among strongly downregulated miRNAs in the DJ-1 overexpression setting. Of particular interest is AGO1 (Argonaute 1), a core component of the miRNA machinery whose functions extend beyond canonical cytoplasmic post-transcriptional regulation. Previous study that qualified the broad effect of silencing AGO-1 by siRNA in HEK293 revealed a modest impact on target suppression (Schmitter et al, 2006). AGO1 has also been shown to interact with RNA polymerase II and associate with promoters of actively transcribed genes, suggesting a role in transcriptional regulation (Huang et al, 2013). Thus, the predicted contribution of AGO1 by the radiation-responsive miRNA population may influence both post-transcriptional and transcription-associated processes and calls for further inspection. Although somewhat speculative, AGO1 remodeling could contribute to the redistribution of miRNA-mediated regulation during the cellular response to DNA damage. This possibility is particularly relevant in DJ-1-OX cells, where irradiation produced extensive remodeling of the miRNA landscape and enrichment of affected targets in DDR and cancer-associated pathways.

A further layer of complexity emerged from the comparison between miRNA precursor transcripts and their corresponding mature products in the DJ-1 OX setting (Fig. 6). The mechanism of miRNAs action in attenuating translation and consequently downregulated targeted transcripts is the outcome of upstream layer of miRNA transcription (reflected by precursor abundance), pre-miRNA successful processing and strand selection (i.e., conversion of precursor to mature miRNA). Then, a layer of stability and turnover that calls for balancing RISC and accessory factor and only then the functional engagement can be achieved (i.e., AGO/RISC binds to miRNA binding sites for target repression). Our data is consistent with the first layers and not explicitly with the functional engagement. Therefore, in future work we will inspect not only the miRNA machinery (Table 1) but the mechanisms responsible for stabilized or reduced miRNA expression (Kai & Pasquinelli, 2010).

The data indicate that irradiation under elevated DJ-1 levels engages both transcriptional and post-transcriptional modes of miRNA regulation. Among upregulated miRNAs, mature accumulation frequently exceeded detectable precursor induction, pointing to a substantial contribution of processing or stability mechanisms. Conversely, downregulated miRNAs showed only partial concordance between precursor and mature levels, consistent with combined effects of transcriptional repression, cellular redistribution or altered maturation dynamics (van Wijk et al, 2022). Together, these observations reinforce the notion that DJ-1 overexpression does not merely alter miRNA abundance but is associated with reconfiguration of miRNA turnover during genotoxic stress adaptation.

Conceptually, genotoxic stress represents a highly dynamic perturbation that is tightly regulated in time and space. The DDR enables rapid yet reversible adaptation through checkpoint activation and repair pathways, thereby preserving genome integrity while preventing premature engagement of irreversible cell fate decisions such as apoptosis (Dufey et al, 2020). miRNAs are uniquely suited for this role: they respond rapidly, do not require the resource and energy demands of protein synthesis, can be redistributed intracellularly and extracellularly, and by utilizing modest cell resources can regulate their own processing and stability. Our findings position DJ-1 dependent miRNA remodeling as a mechanism of fast translational buffering that promotes adaptive recovery. In a future work we will apply AGO-CLIP and small-RNA immunoprecipitation after X-ray to monitor cellular redistribution of AGO-loaded miRNAs and the kinetics of P-bodies along cells recovery and adaptive response.

Several limitations should be considered in this study. First, our conclusions rely primarily on comparative expression and the solid testing of controls (e.g., untreated cells, empty plasmid, siRNA RULC). While, we rely of our analysis of experimental evidence from miRTarBase, a direct validation of key miRNA-target interactions and functional assays of translational output are needed. Second, although in major cell lines, the steady state level of DJ-1 is substantial, our experiments were performed in HEK293 cells with an increase in DJ-1 that are beyond physiological range. It is likely that testing DJ-1 dependent miRNA plasticity in cells that is naturally under oxidative and genotoxic pressure such as neuronal dopaminergic cells and tumor models could mimics disease state such as ALS, Parkinson disease (Weng et al, 2023) and cancer metastasis. Third, while the impact of DJ-1 cellular manipulation on the protein and other lncRNAs were established (Zohar et al, 2025), the analyses addressed cells as a whole without implicitly address the redistribution of miRNAs among compartments (mitochondria, nucleus, cytosol, exosomes). Lastly, because all measurements were obtained at a single early post-irradiation time point (6 h), the study cannot distinguish transient remodeling from sustained changes in miRNA biogenesis and turnover. Despite these limitations, our results have potential implications for further studies and to other cellular systems. DJ-1 is frequently is elevated in metastatic cancers and has been implicated in stress tolerance and therapy resistance. The DJ-1-miRNA axis described here may contribute to radiation resistance by enhancing post-transcriptional buffering capacity. Modulating DJ-1 levels could therefore represent strategies to sensitize tumors to genotoxic therapies (Olivo et al, 2022). Conversely, in neurodegenerative contexts where DJ-1 function is compromised, impaired miRNA buffering may exacerbate vulnerability to oxidative or DNA damage.

Taken together, our data support a model in which high DJ-1 levels enable coordinated miRNA remodeling that halt and stabilizes translation and metabolism, promoting adaptive resilience. Low DJ-1 levels, in contrast, are associated with reduced regulatory plasticity and reliance on less coordinated stress pathways and loss of homeostasis. DJ-1 thus emerges as an integrator linking DNA damage sensing to miRNA-dependent translational control, with implications for cancer, aging, and neurodegeneration.

## Author Contributions

K.Z. led the project from the experimental design, RNA-seq analysis, data analysis, visualization. M.L. served in mentoring, visualization and wrote the initial manuscript. The authors have reviewed and edited the output and take full responsibility for the content of this publication.

## Institutional Review Board Statement

Not applicable.

## Informed Consent Statement

Not applicable.

## Data Availability Statement

The raw data was submitted to ArrayExpress Accession E-MTAB-14761. Detailed summary of the quality control and reads assignment is provided in Supplementary Tables. The source data for the figures are available in Supplementary Tables S1-S5.

## Acknowledgments

We thank the members of the Linial’s lab for useful suggestions and fruitful discussions. We thank Tsiona Eliyahu for preparation of total RNA used in this study. We thank the Genomic and Sequencing Center of the Hebrew University for their support. We thank the Clore Israel Foundation for their Scholar program and the fellowship granted to K.Z.

## Abbreviations

AD: Alzheimer’s disease
BSA: Bovine serum albumin
CNS: Central nervous system
DEG: Differentially expressed gene
DMEM: Dulbecco’s modified Eagle medium
FC: Fold change
FDR: False discovery rate
HPA: Human proteome atlas
GO: Gene ontology
IFN: Interferon
NT: Not treated
PCA: Principal component analysis
PCR: Polymerase chain reaction
PD: Parkinson’s disease
RNA-seq: RNA sequencing analysis
RT: Reverse transcription
TMM: Trimmed mean of means

